# Dysregulation of cell migration by matrix metalloproteinases in geleophysic dysplasia

**DOI:** 10.1101/2025.01.30.635317

**Authors:** Alejo A. Morales, Vladimir Camarena, LéShon Peart, Sarah Smithson, Lindsay Shaw, Lucy Webber, Jose M. Negron, Juan E. Sola, Ann-Christina Brady, Katherina Walz, Gaofeng Wang, Mustafa Tekin

**Author notes:** Address correspondence to: Mustafa Tekin, John T. Macdonald Foundation Department of Human Genetics, University of Miami Miller School of Medicine, 1501 NW 10th Avenue, BRB-610 (M-860), Miami, Florida 33136, USA.

## Abstract

Geleophysic dysplasia (GD) is characterized by short stature, brachydactyly, joint limitations, a distinctive facial appearance, as well as cardiac and respiratory dysfunction that can be life-threatening. GD is caused by pathogenic variants in the *ADAMTSL2*, *FBN1,* or *LTBP3* genes. While dermal fibroblasts derived from affected individuals have shown poor organization of the extracellular matrix (ECM), it remains elusive how the disorganized ECM contributes to GD pathogenesis. To understand the molecular mechanisms in GD, we isolated and characterized primary human dermal fibroblasts from affected individuals with *ADAMTSL2* and *FBN1* variants. We found that the secretion of ECM proteins including ADAMTSL2, FBN1, and Fibronectin were impaired in GD fibroblasts. Increased cell migration was observed in GD fibroblasts carrying *ADAMTSL2* or *FBN1* variants, which was associated with up-regulation of MMP-1 and MMP-14, two proteases related to cell mobility. The enhanced cell migration and up-regulation of MMP-1 and MMP-14 were corroborated in mouse primary dermal fibroblasts carrying pathogenic variants in *Adamtsl2* and in lung and heart tissues from *Adamtsl2*-*knockout* mice. A pan MMP inhibitor, GM6001, inhibited the migration of GD fibroblasts. Overall, our results suggest that MMP-1/-14 up-regulation play a role in the development of GD and may be utilized as a treatment target.

## Introduction

Geleophysic dysplasia (GD) is a rare and progressive genetic condition characterized by short stature, brachydactyly, joint limitations, and a unique facial appearance as well as cardiac and respiratory dysfunction that result in poor prognosis [1–7]. Three genes have been associated with GD, resulting in distinct forms of the disorder. Geleophysic dysplasia type-1 (GD1, GPHYSD1, OMIM 231050) is an autosomal recessive disorder caused by *ADAMTSL2* variants (OMIM 612277) [8]. Geleophysic dysplasia type-2 (GD2, GPHYSD2, OMIM 614185) is an autosomal dominant disorder caused by variants in exon 41 or 42 of *FBN1* (OMIM 134797) [9]. Geleophysic dysplasia type-3 (GD3, GPHYSD3, OMIM 614185) is an autosomal dominant disorder caused by *LTBP3* variants (OMIM 602090) [10]. Variants in *ADAMTSL2* and *FBN1* can lead to their impaired secretion [4,11,12]. Multiple studies have shown that the poor secretion of these proteins affects the extracellular matrix (ECM) organization; however, the underlying molecular mechanism has not yet been clarified.

ADAMTSL2 is a secreted matricellular glycoprotein and a member of the A Disintegrin-like And Metalloprotease with Thrombospondin motifs (ADAMTS) superfamily, which lacks the metalloprotease domain, a characteristic of this family [13–15]. While ADAMTSL2 function is not well-characterized, recent reports have suggested an important role during the fibrillin microfibril organization [16–19], and as a regulator of the Transforming Growth Factor Beta (TGF-β) availability [8,9,20,21]. These findings can be explained, in part, due to the interaction of ADAMTSL2 with other proteins in the extracellular space, including FBN1 and LTBP1 [8,20,21].

Fribrillin-1 is a large calcium binding extracellular matrix glycoprotein and a major element of the microfibrils in the extracellular space, which plays multiple roles in the elastic and nonelastic fibers in the connective tissue [22–24]. It is a multidomain protein considered also a regulator of the bioavailability of multiple molecules, most notably TGF-β [25,26]. Marfan syndrome is the best known fibrillinopathy [27,28], but some other pathogenic variants in *FBN1* have been also reported in other disorders, including GD2 and Weill-Marchesani syndrome (WMS) [28]. Specifically, mutations in exons 41-42, which encode the TGF-β-binding protein-like domain 5 (TB5) of FBN1, have been proposed, together with the dysregulation of the TGF-β signaling, as the molecular mechanisms for GD2 [8,9].

Both ADAMTSL2 and Fribrillin-1 potentially interact with other ECM proteins, such as matrix metalloproteinases (MMPs). MMPs are a large family of Ca^2+^-dependent Zn^2+^-containing endopeptidase enzymes, classified into multiple and different groups, based on their primary substrate specificity or their cellular localization: collagenases (MMP-1, -8, and -13), gelatinases (MMP-2 and -9), stromelysins (MMP- 3, -10, and -11), matrilysins (MMP-7 and -26), macrophage elastase (MMP-12), membrane-type MMPs (MMP-14, -15, -16, -17, -24, and -25), and others (MMP-19, -20 -21, -23A/B, -27, and -28) [29–31]. They process and degrade major components of the ECM during tissue remodeling, affecting multiple cellular processes such as proliferation, inflammation, migration, and survival [32–35]. MMPs are produced as inactive zymogens, which require proteolytic activity for the active form. MMP activity is finally controlled by their four natural inhibitors or tissue inhibitors of metalloproteinases (TIMPs) [36,37]. Together, they have numerous roles in physiological and pathological conditions, including cancer, cardiovascular diseases, fibrotic disorders, lung diseases, and inflammation [38,39].

The molecular mechanisms underpinning GD remain largely elusive. In this study, we characterized dermal fibroblasts from individuals with GD caused by *ADAMTSL2* or *FBN1* variants and compared them with control fibroblasts. Our findings demonstrate that patient fibroblasts have an enhanced cell migration, mediated by an overall increase of MMPs secretion. Among MMPs we found that the patient fibroblasts had increased levels of MMP-1 and MMP-14, two enzymes with critical roles in cell migration and in the ECM protein degradation and remodeling. This is the first report to identify up-regulation of MMPs in fibroblasts from GD suggesting a possible role of these proteases in the molecular mechanisms for the disease.

## Results

### Establishment of fibroblasts from patients with GD1 and GD2

We and others have previously shown that mutations in *ADAMTSL2* and *FBN1* genes resulted in their impaired secretion [8,9,12,40]. To elucidate the impact of variants in *ADAMTSL2* and *FBN1* genes in cellular and molecular processes, we characterized primary human dermal fibroblasts from ADAMTSL2- related GD (GD1) and FBN1-related GD (GD2) and compared them with fibroblasts from asymptomatic individuals as controls. The clinical diagnosis and variants identified in all the patients included in this study are shown in Table 1. GD1 patients are compound heterozygous for *ADAMTSL2* variants, while GD2 patients are heterozygous for *FBN1* variants. Since LTBP3-related GD (GD3) represents less than 1% of total affected GD patients [1], they have not been included in our study. By the time we obtained the skin biopsy to isolate the dermal fibroblasts, all patients showed the clinical characteristics of GD, including short stature and cardiorespiratory conditions. All patients and healthy volunteers were evaluated at the University of Miami or by collaborators, and all diagnoses were confirmed by sequencing as *ADAMTSL2* or *FBN1* gene variants.

**Table 1.** Phenotypic and genotypic findings of the study cohort.

| Patient ID | HDFn | GD016E | GD017E | GD001 | GD006 | GD004 | GD009 | GD012 |
| --- | --- | --- | --- | --- | --- | --- | --- | --- |
| Diagnosis | CT | CT | CT | GD1 | GD1 | GD2 | GD2 | GD2 |
| Sample ID | CT | CT-1 | CT-2 | GD1-1 | GD1-2 | GD2-1 | GD2-2 | GD2-3 |
| Height |  |  |  | 86.4 cm<br>(z-score-1.16) | 120 cm<br>(z-score-2.21) | extremely<br>short (z - N/A) | 117 cm<br>(z-score-2.53) | 95.5 cm<br>(z-score-7.5) |
| Cardiac pathology |  |  |  | PS, AS, PDA, VSD | AS, PS, MS | AS, MS, PS, HF | normal | normal |
| Respiratory involvement |  |  |  | Frequent<br>infections | Frequent<br>infections,<br>tracheostomy | Frequent<br>infections,<br>tracheostomy | Frequent<br>infections | Frequent<br>infections,<br>tracheostomy |
| Mutations |  |  |  | **ADAMTSL2**<br>c.182G>A<br>**p.R61H**<br>c.493G>A<br>**p.A165T**<br>compound<br>heterozygous | **ADAMTSL2**<br>c.542T>C<br>**p.V181A**<br>c.707C>T<br>**p.P236L**<br>compound<br>heterozygous | **FBN1**<br>c.5284G>A<br>**p.G1762S**<br>heterozygous | **FBN1**<br>c.5183C>T<br>**p.A1728V**<br>heterozygous | **FBN1**<br>c.5243G>C<br>**p.C1748S**<br>heterozygous |
| Sex | male | male | male | male | female | female | male | male |
| Skin biopsy (location) | foreskin | foreskin | foreskin | shoulder | unknown | shoulder | shoulder | shoulder |
| Age - years (months) | neonate | 0 (1) | 0 (6) | 1 (11) | 12 (0) | 28 (9) | 8 (8) | 11 (0) |

Asymptomatic Control (CT), Geleophysic Dysplasia type-1 (GD1) and Geleophysic Dysplasia type-2 (GD2). Pulmonary stenosis (PS), aortic stenosis (AS), patent ductus arteriosus (PDA), vasospastic angina (VSA), mitral stenosis (MS) and heart failure (HF).

No significant morphological difference was found between mutant and control fibroblasts. All presented a similar spindle-shaped morphology with a big oval nucleus, multiple big vacuoles in the cytoplasm, and similar size with an average of 20 µm as detached cells. After confluency, all fibroblast cultures showed the characteristic parallel array arrangement and swirling pattern, although for patient cells a delay in the time to reach this pattern was observed. Since fibroblast swirling in culture is in part due to their elongated shape, proliferation, and motility [41], we next studied these cell properties. Patient fibroblasts did not show any other significant morphological differences compared to the control cells under the light microscope.

### GD fibroblasts migrate faster than controls

To further characterize the fibroblasts, we determined their proliferation for 10 days, in a 96-well plate setting using a lactate dehydrogenase (LDH) assay. We determined the proliferation under two initial cell densities: 1) A low cell density with 1.5×10^4^ cells/well to better study their proliferation, and 2) A high cell density with 7.5×10^4^ cells/well, to study their proliferation with the required confluency to produce mature ECM. As expected, lower initial cell density showed a faster rate of proliferation for all fibroblasts as determined by the LDH assay, mainly after 7 days (Figure 1A, B). The total LDH in 10 days, as an indicator of the total amount of cells, increased an average of 4.2 times for the lower initial cell count and only 1.7 times for the higher initial cell count. Although fibroblasts individually proliferated at different rates, overall, they did not show any significant difference in proliferation compared to the control cells. Similar results were obtained when cells were confluently seeded to maximize ECM production (Figure 1B). This is particularly important as we next checked the fibroblast migration, and these results exclude any significant contribution of proliferation to cell migration. The high cell density protocol was used for all the following experiments.

**Figure 1.**
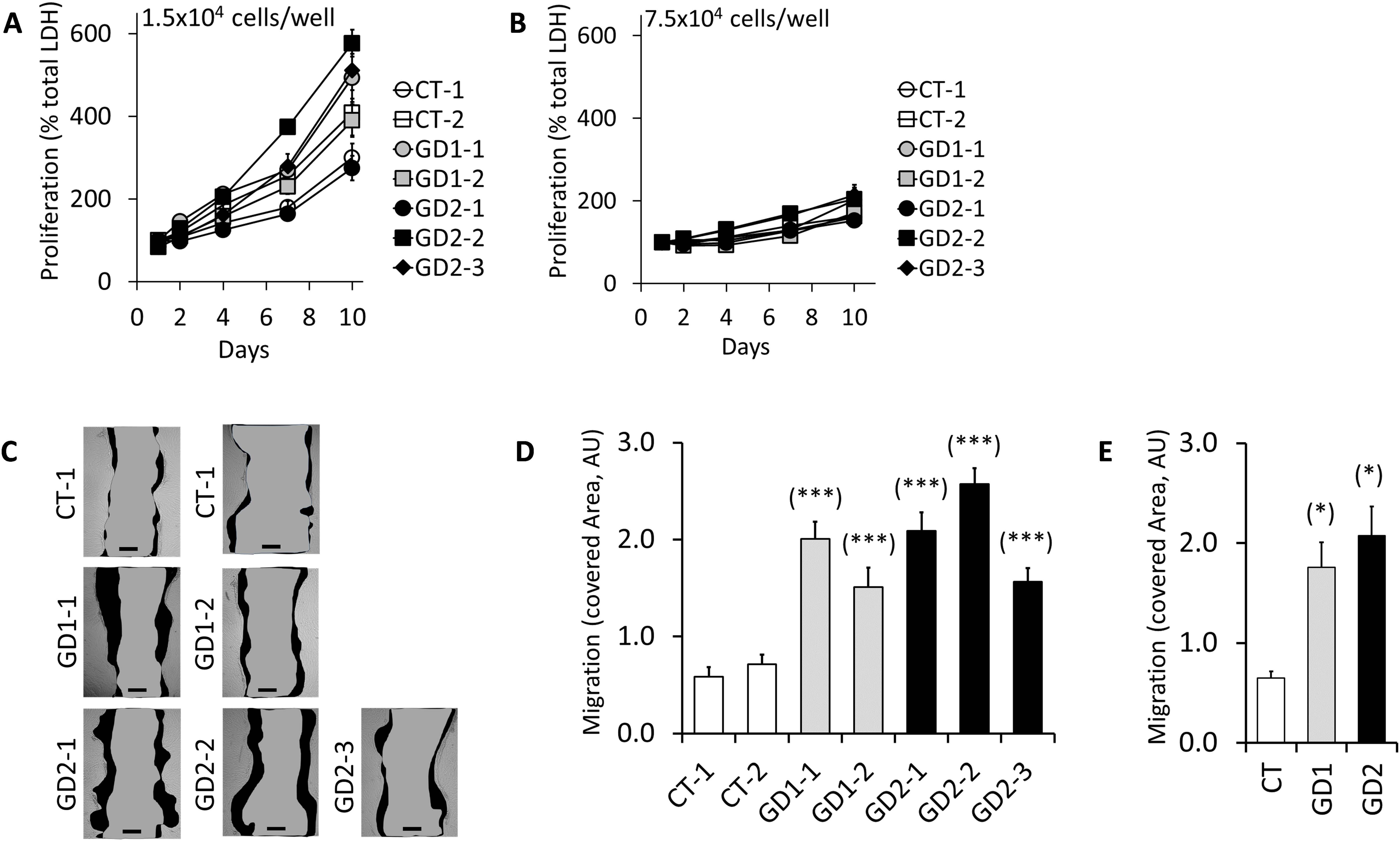
Patient fibroblasts migrate faster than control fibroblasts even without significant difference in cell proliferation. **(A)** Cell proliferation was assessed using the LDH assay and two different initial densities: 1.5×10^4^ cells/well and **(B)** 7.5×10^4^ cells/well. Averages were calculated for 3 independent experiments. No significant difference was found between the cells when they were compared to the CT fibroblasts. **(C)** Cell migration was assessed using the wound healing scratch assay for 8 hours. Scale bar = 50 µm. Representative images were selected from five independent experiments. **(D)** Covered area (dark, in Arbitrary Units) was calculated using ImageJ software. **(E)** Average for CT (2 primary cell samples), GD1 (2 samples) and GD2 (3 samples) were calculated and compared. Significance was measured using a t-test and values were compared versus CT fibroblasts (* p<0.05 and ***p<0.001).

To assess the fibroblast migration, we used the two-dimensional wound healing assay (scratch migration assay or simply scratch assay). After the scratch, the cell migration was calculated based on the covered scratch area for 8 and 24 hours. At 24 hours most of the scratches for patient cells were closed and the data was excluded (data not shown). Furthermore, as fibroblasts showed no significant difference in proliferation for 10 days, and no proliferation for the first 48 hours (Figure 1B), the scratch closure after only 8 hours was mainly caused by cell migration. Although we observed some variations between the samples, all patient fibroblasts, individually, migrated statistically faster than the control cells, covering a higher scratched area at 8 hours (Figure 1C, D). We also performed the cell migration analysis by clinical groups, and we determined that GD1 and GD2 fibroblasts migrated 2.7 and 3.2-times faster than the control group, respectively (Figure 1E). The migration of GD1 and GD2 fibroblasts were at the same levels (p=0.5042). Overall, our data suggest that GD fibroblasts migrate faster than the controls.

### Secretion of ADAMTSL2, FBN1, and Fibronectin is impaired in GD fibroblasts

Since the rate of cell migration adapts to physical cues in the extracellular environment, we set up to investigate the ECM in GD fibroblasts. First, we studied the presence of mutant proteins into the ECM. Previous reports have shown that mature ECM is produced after 4-7 days of fibroblasts culture, with greater ECM accumulation at 7 days [20]. We obtained and analyzed three protein lysates: intracellular (IC) lysate, ECM lysates, and total lysates (Total), which should contain all the IC and ECM proteins, and serve as a control for the individual lysates, especially the technically challenging ECM lysates.

ADAMTSL2 was detected mainly as an IC protein (band-1 and band-2 in Figure 1S), with very low detection in the ECM, when it was compared with the IC levels. Fibronectin and GAPDH were used as controls for the extracellular and intracellular fractions respectively. FBN1 and fibronectin proteins were mainly detected in the ECM with very low expression in the IC compartment (Figure 1S).

All patient fibroblasts showed an impaired protein deposition of ECM components: ADAMTSL2, FBN1, and fibronectin. As previously reported [40], ADAMTSL2 mutations in GD1 fibroblasts did not affect the intracellular protein expression, and all cells showed similar levels of expression, compared to the control cells (Figure 2A, B). However, ADAMTSL2 mutations affected the accumulation of the protein in the ECM (arrow, the most abundant band in Figure 2A). A drastic reduction of ADAMTSL2 protein was observed in GD1 patient fibroblasts. Unexpectedly, GD2 fibroblasts, with *FBN1* mutations, also failed to incorporate ADAMTSL2 protein into the ECM, even with normal intracellular expression (Figure 2A, B).

**Figure 2.**
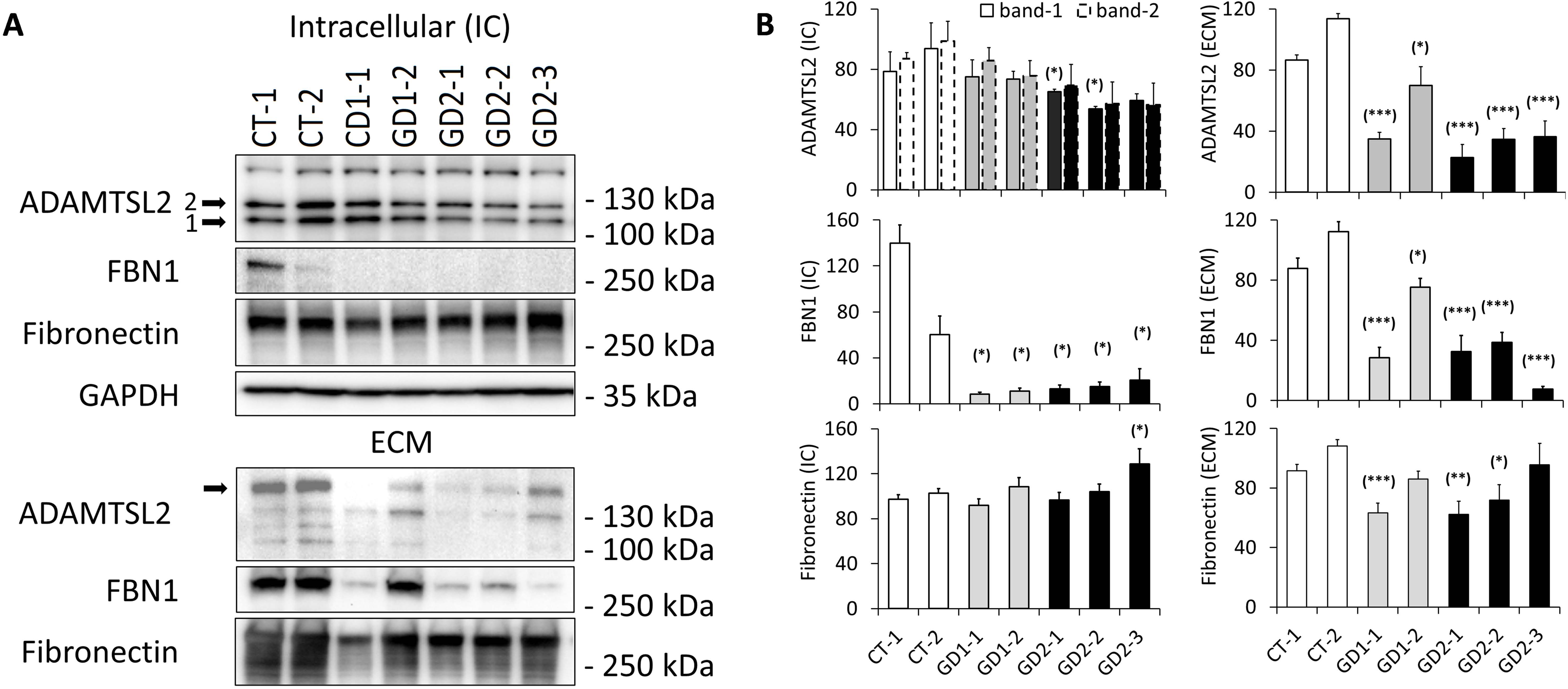
Patient fibroblasts show impaired ECM protein accumulation. Cell lysates were prepared for all fibroblasts culture; and intracellular lysates (IC) and extracellular matrix lysates (ECM) were analyzed by WB, using GAPDH and Fibronectin as IC and ECM controls, respectively. ADAMTSL2 was detected mainly in the IC compartment (band-1 and band-2) while FBN1 and Fibronectin proteins were detected mainly in the ECM space. **(A)** ADAMTSL2, FBN1 and fibronectin deposition in the ECM were impaired in patient’s fibroblasts. Intracellular FBN1 was decreased also in patient’s cells. Representative images were selected from four independent experiments. **(B)** Densitometry analysis was completed using ImageJ software. Significance was measured using a t-test and values were compared versus CT fibroblasts (* p<0.05, ** p<0.01, and *** p<0.001).

*FBN1* mutations in GD2 cells affected the intracellular expression of the FBN1 protein as well as the accumulation in the ECM. Surprisingly, GD1 fibroblast, with WT variants for *FBN1* but *ADAMTSL2* mutations, failed to incorporate FBN1 protein in the ECM (Figure 2A, B). Compared to the CT fibroblasts, most of the patient cells failed to accumulate FBN1 in the ECM.

*ADAMTSL2* and *FBN1* mutations did not affect the synthesis of fibronectin in GD1 and GD2 cells, respectively; but its accumulation in the ECM was decreased in most patient fibroblasts (Figure 2A, B).

Patient fibroblasts generally showed poor ECM accumulation and organization (supplemental Figure 2S). They failed to form the parallel lines of fibers characteristic in control cells and multiple empty areas were present in the ECM from patient fibroblasts. There was an interesting correlation between the reduced ECM in the patient fibroblasts with their delay in the swirling pattern.

Overall, the ECM and the secretion of ADAMTSL2, FBN1, and Fibronectin is impaired in GD fibroblasts. We subsequently verified if the reduced ECM deposition could be underpinning the increased migration observed in patient fibroblasts (Figure 1C-E).

### ECM from healthy fibroblast does not normalize the migration of GD fibroblasts

To check if the enhanced migration observed in patient fibroblasts was promoted by a weak or reduced ECM, we isolated an abundant ECM from a control fibroblast cell line and grew the cells on top. For this experiment, we selected two control cell lines and two GD1 fibroblasts (Figure 3A-C). The isolated ECM from control cells (CT), showed the highest expression of ADAMTSL2, FBN1, and fibronectin from all the control cells used in our studies (data not shown). There were no significant changes in the cell growth for any of the fibroblasts studied under these conditions. Surprisingly, the two control cells slightly increased their migration, although no statistical significance was obtained. Results from the patient’s fibroblasts were not consistent. One of the patient fibroblasts (GD1-2) showed a reduced cell migration when it was grown on healthy derived-ECM, although no statistical significance was obtained. Interestingly, this patient fibroblast showed the most abundant ECM accumulation along the GD groups (Figure 2 and Figure 2S). Results for the other GD1 patient fibroblasts showed no effect on cell migration when cells were exposed to healthy derived-ECM (GD1-1 in Figure 3C). Taken together, this experiment suggests that exogenous ECM, despite derived from a healthy cell culture, is not enough to rescue the enhanced migration observed in all patient fibroblasts.

**Figure 3.**
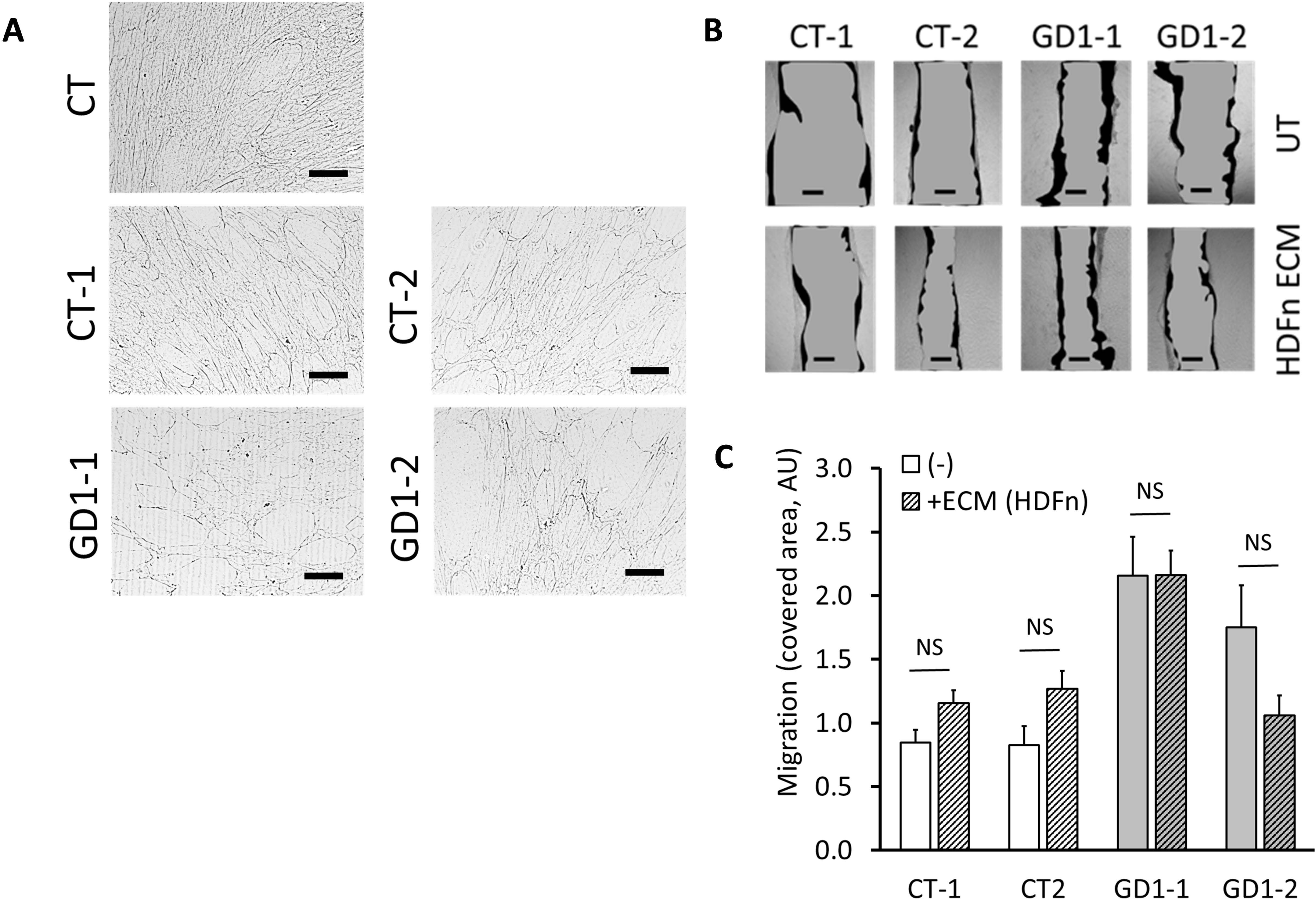
ECM from control fibroblasts does not inhibit migration of patient fibroblasts. **(A)** ECM deposition for GD1 patient fibroblasts was impaired when compared with CT fibroblasts. Scale bar = 200 µm. **(B)** Cells were grown on top of the ECM from control fibroblast (HDFn) for 48 hours (confluency). Cell migration was assessed using the wound healing scratch assay for 8 hours. Scale bar = 50 µm. Representative images were selected from four independent experiments, using two CT fibroblasts and two GD1 patient’s fibroblasts. **(C)** Covered area (dark, in Arbitrary Units) was calculated using ImageJ software. Significance was measured using a t-test and values were compared versus CT fibroblasts (NS, not significant).

### The expression of MMPs is upregulated in GD fibroblasts

We further expanded our analysis and compared the RNA-Seq transcriptome data for all fibroblasts included in this study. Initial unsupervised hierarchical clustering of top differential transcripts showed a clear separation of the two main groups: controls vs GD cells (data not shown); but there was not an evident explanation for the enhanced migration of patient fibroblasts. We focused then on genes that could affect the ECM and identified that MMP genes were differentially up-regulated in the patient fibroblasts. The effect of MMPs on cell migration has been previously described [42–45]. We performed a supervised analysis with all MMPs included in our data. While most of the MMPs were up-regulated and clustered together in the patient fibroblasts; the four natural MMP inhibitors (TIPM1-4), also clustered together but were down-regulated. Among the up-regulated MMPs found in patient fibroblasts, MMP1, MMP3, MMP8, MMP12, MMP14, and MMP27 showed the strongest increase (Figure 4A). It is also important to point out that the ratio of MMPs to TIMPs strongly indicated a higher MMP activity in patient fibroblasts.

**Figure 4.**
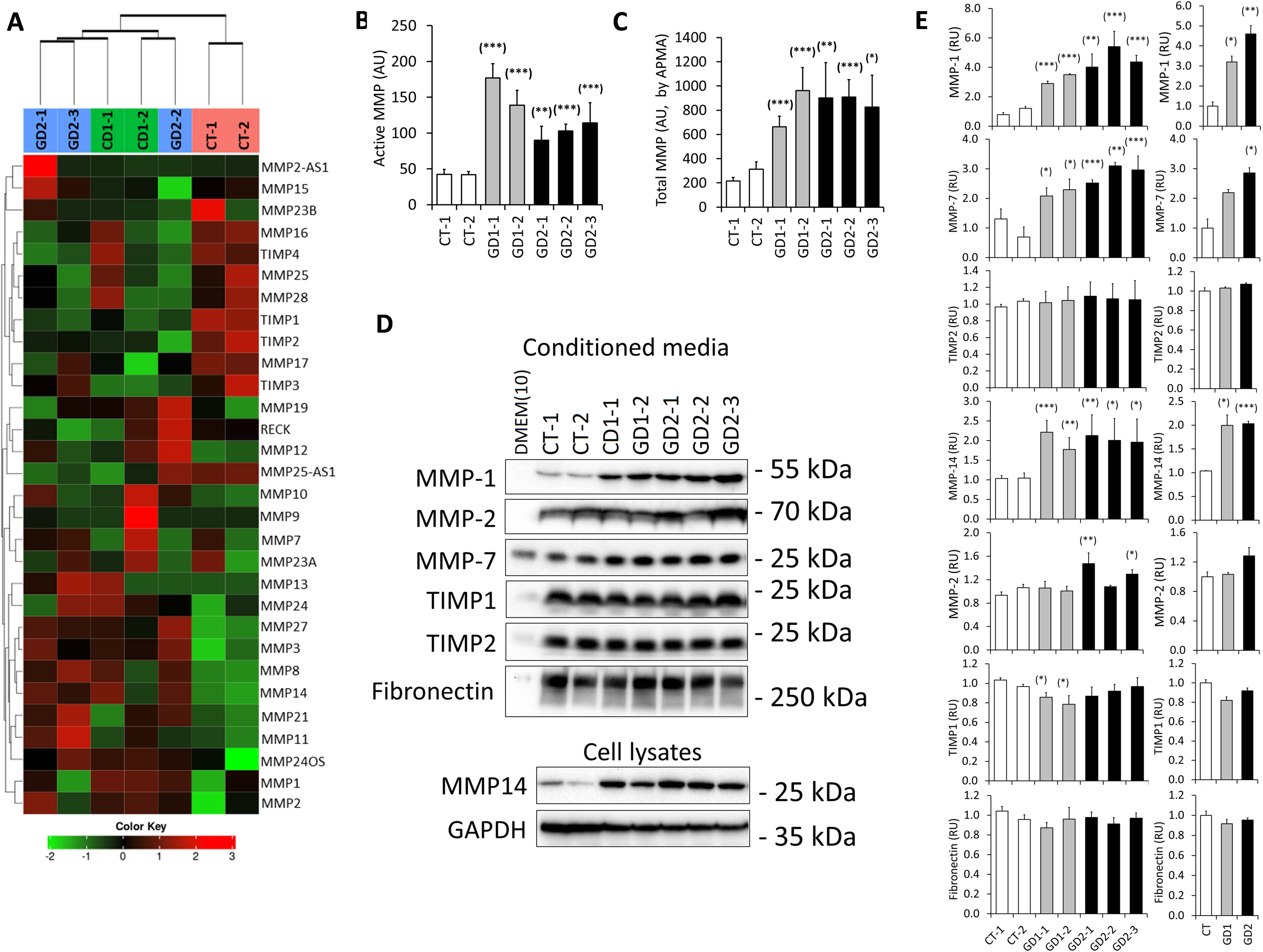
Patient fibroblasts express high levels of MMPs. **(A)** RNA-Seq transcriptome analysis of patient fibroblast from GD1 (N=2) and GD2 (N=3) compared to controls (N=2). Semi-supervised hierarchical clustering of MMPs and TIMPs transcripts. **(B)** Cells were grown, and medium was not changed for 10 days. Instead, small aliquots of fresh medium were added every other day. Active MMPs were determined in the conditioned medium using a fluorometric assay (MMP Activity Assay Kit). Medium (DMEM+10% FBS) was used as negative control. The fluorescence signal, as Relative Fluorescence Unit (RFU), was averaged for each sample from five independent experiments. All samples were compared to the CT average. **(C)** Conditioned media were treated with p-Amino phenyl mercuric acetate (APMA) for two hours at 37°C; and Total MMP activity was determined following a similar protocol. **(D)** MMP proteins were analyzed by WB using the CM (secreted MMPs) or cell lysates (MMP14, a transmembrane protein). Medium [DMEM(10), DMEM+10% FBS] was used as negative control. GAPDH was used as loading controls for CM and cell lysates, respectively. Representative images were selected from four independent experiments. **(E)** Densitometry analysis was performed using ImageJ software. Significance was measured using a t-test and values were compared versus CT fibroblasts (* p<0.05, ** p<0.01, and *** p<0.001).

We next studied the MMP protein profile for all fibroblasts and selected a green fluorometric assay to determine the overall MMP activity in the cells. This assay uses a fluorescence resonance energy transfer (FRET) peptide as an indicator, the peptides are quenched to each other but MMP cleavage results in dequenching and increased fluorescence. We used conditioned media that were not changed for 10 days. Instead, small aliquots of fresh media were added every other day, and MMPs accumulated in the conditioned media. Figure 4B shows that the overall MMP activity was significantly higher in conditioned media from all patient fibroblasts when compared with control cells, with an average of 3.0-fold change for all GD cells. As this measurement represents the portion of all active MMPs, we wanted to determine also the total amount of MMP present in the conditioned media. For this purpose, we used 4- aminophenylmercuric acetate (APMA), an organomercurial that activates most of the MMPs (46). We treated all conditioned media with 2 mM APMA at 37°C for 2 hours. Similar to the active MMP activity, the total MMP activity was significantly higher in conditioned media from patient fibroblasts (Figure 4C), with an average of 3.2-fold increase, when compared with controls. These results suggest that there is a higher MMP activity and production in patient conditioned media.

As some MMPs are not secreted but are attached to the cell membrane, we next selected some of the most up-regulated MMPs genes, to check their protein expression by WB (Western blotting). For secreted MMPs we used conditioned media collected after 10 days. Membrane-associated MMP-14 (MT1-MMP) was determined in cell lysates (RIPA buffer lysates). Three MMPs were strongly upregulated in patient fibroblasts: MMP-1, MMP-7, and MMP-14 (Figure 4D, E).

MMP-1 was not detected in the cell culture control media [DMEM(10), DMEM with 10% FBS] kept in parallel wells without cells. When compared to control cells, all patient fibroblasts strongly up-regulated the MMP-1 secretion into the conditioned media, ranging from 3 to 5.5-fold changes. Based on the RNA-Seq results, most of the patient fibroblasts also showed up-regulation in the MMP1 gene, suggesting an increased transcription of the gene as the mechanism for the MMP-1 up-regulation.

MMP-7 protein was detected in the DMEM(10), which suggests a species cross-reactivity of the antibody, detecting bovine MMP-7 from the FBS. Nevertheless, we also detected the MMP-7 secretion in all conditioned media from fibroblasts; and it was upregulated in the patient cells.

Membrane-associated MMP-14 was also up-regulated in the patient cell lysates, with an average of 2-fold changes (Figure 4D, E). Similar to the MMP-1, the protein up-regulation correlated with the MMP-14 gene up-regulation (Figure 4D, E). All this together, strongly suggested an up-regulation mechanism at the gene transcription level, not only for MMP-1 but also for MMP-14.

Detection of some MMPs by WB was challenging, as they have a molecular weight similar to the BSA present in the conditioned media (FBS). To confirm the protein expression, we also performed an antibody array, that included seven MMPs and three TIMPs. We confirmed the up-regulation of multiple MMPs such as MMP-1, MMP-2, MMP-3, MMP-8 and MMP-10, while TIMP4 was down-regulated for the GD1 conditioned medium when compared to the control cells (Figure 3S). Taking together, these results suggest that MMPs are up-regulated in GD fibroblasts including MMP-1/-14.

### Validation of enhanced cell migration and up-regulation of MMP-1/-14 in GD1 mouse fibroblasts

We recently reported the development and characterization of a mouse model for GD, which replicates the genetic variants of the GD1-1 patient who is compound heterozygous for *ADAMTSL2* p.R61H and p.A165T variants [40]. We next tested if our findings in human dermal fibroblasts could be replicated in the mouse model. Using the explant protocol, we isolated mouse dermal fibroblasts from two adult WT mice and two adult *ADAMTSL2*-p.R61H-p.A165T mice (RA or R61H/A165T) and we checked the cell migration and the MMP profile for these cells. WB showed multiple non-specific bands and a band (probably a post translational modification form of ADAMTSL2) was reduced in the R61H/A165T lysates (Figure 5A) [40]. Post translational modification like N-glycosylation and O-fucosylation have been reported as essential steps for ADAMTSL2 secretion [15,18]. The murine R61H/A165T fibroblasts also showed a strong and significantly enhanced migration compared to the WT controls (Figure 5B, C), confirming the migration phenotype in a different species. Interestingly, both MMP-1 in the 10-day conditioned media and MMP-14 in the cell lysates were also significantly up-regulated in the R61H/A165T fibroblasts (Figure 5D, E), thus confirming the upregulation of cell migration and MMP-1/-14 in GD. All together these results suggest that blocking the MMPs activity might normalize the enhanced GD migration.

**Figure 5.**
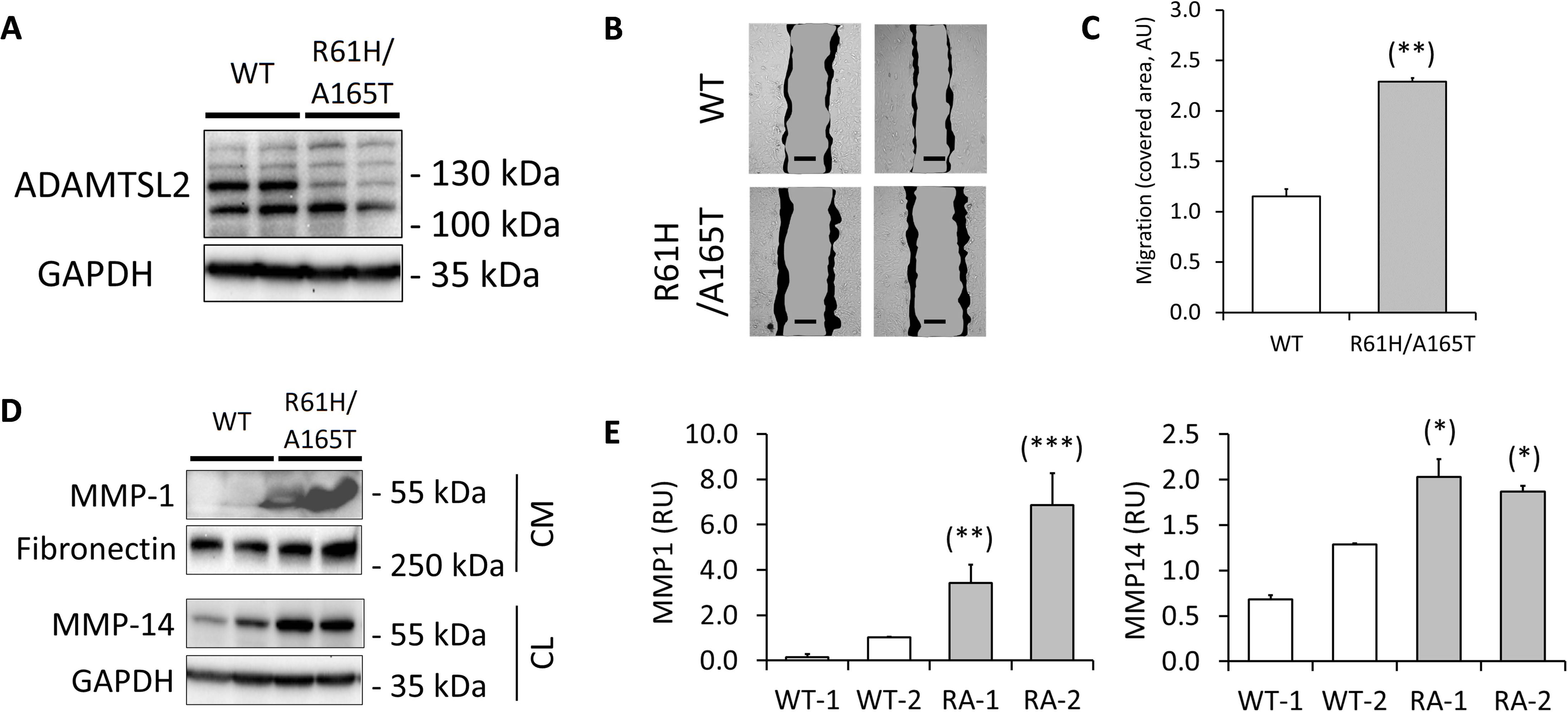
Mouse dermal fibroblasts from GD1 mouse model show enhanced migration and up- regulation of MMP1 and MMP14. **(A)** WB showed multiple non-specific bands for fibroblasts lysates and a band (most probably attributed to the N-glycosylated form of ADAMTSL2) was strongly reduced in the R61H/A165T lysates. **(B)** Cells were grown for 48 hours (confluency). Cell migration was assessed using the wound healing scratch assay for 8 hours. Scale bar = 50 µm. Representative images were selected from three independent experiments, using two WT fibroblasts and two R61H/A165T fibroblasts. **(C)** Covered area (dark, in Arbitrary Units) was calculated using ImageJ software. **(D)** MMP1 protein was analyzed by WB using 10-days conditioned media and MMP14 was analyzed using cell lysates. GAPDH was used as loading controls for CM and cell lysates. Representative images were selected from four independent experiments. **(E)** Densitometry analysis was performed using ImageJ software. Significance was measured using a t-test and values were compared versus CT fibroblasts (* p<0.05 and ** p<0.01).

### MMPs are upregulated in the lung and heart of ADAMTSL2-KO mice

We next analyzed tissues from *ADAMTSL2-KO* neonate mice, which resemble the Al-Gazali skeletal dysplasia, the most severe form of GD1 with a lethal phenotype [18, 40, 47]. Lung and heart lysates were prepared from tissues obtained during the first 24 hours after birth. Similar to the GD1 mouse fibroblasts (Figure 5A), WB of lung lysates showed multiple non-specific bands while one unique band was absent in the *ADAMTSL2-KO* lysates (Figure 6A). This band corresponded to the same band that was reduced in the R61H/A165T fibroblasts when compared to WT cells. While MMP1 expression showed no significant difference between WT and *ADAMTSL2-KO* lysates, MMP14 was strongly up-regulated in the lungs of *ADAMTSL2-KO* mice (Figure 6A, B).

**Figure 6.**
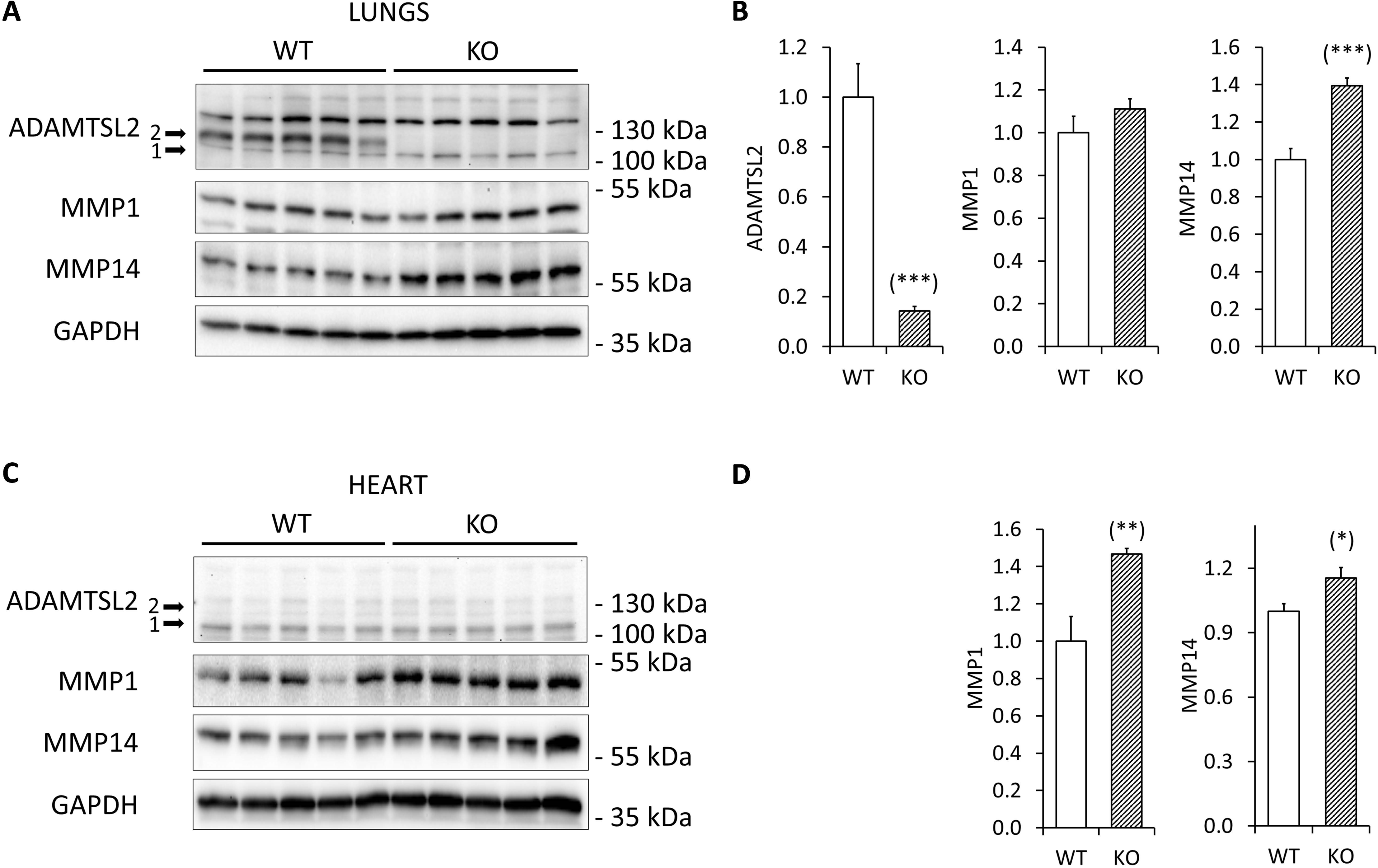
Lung and heart lysates from ADAMTSL2-KO neonate mice showed up-regulation of MMP1 and MMP14. **(A)** WB showed multiple non-specific bands for the WT lung lysates and a band (most probably attributed to the N-glycosylated form of ADAMTSL2) was not present in the ADAMTSL2- KO lysates. Lung lysates from ADAMTSL2-KO mice showed upregulation of MMP14. **(B)** Densitometry analysis for ADAMTSL2, MMP1 and MMP14 was performed using ImageJ software. Significance was measured using a t-test and values were compared versus WT lysates (*** p<0.001). **(C)** WB showed multiple non-specific bands for the WT and ADAMTSL2-KO heart lysates with no significant difference. Heart lysates from ADAMTSL2-KO mice showed upregulation of MMP1 and MMP14. **(D)** Densitometry analysis for MMP1 and MMP14 was performed using ImageJ software. Significance was measured using a t-test and values were compared versus WT lysates (* p<0.05 and ** p<0.01).

Furthermore, MMP1 and MMP14 protein levels were significantly up-regulated in *ADAMTSL2-KO* heart tissues when compared with WT controls (Figure 6C,D). Using Real Time PCR, our earlier work has validated a complete knockdown of *ADAMTSL2* in these animals [40]. However, we were unable to confirm the knockdown of ADAMTSL2 at the protein level in the heart. WB of heart samples showed multiple non-specific bands without obvious differences between WT and *ADAMTSL2-KO* lysates (Figure 6C).

### MMP inhibitor blocks the migration of GD fibroblast

As MMPs have been reported to play important roles in cell migration, we next checked the effect of a general MMP inhibitor on fibroblast migration. We plated the cells, grew them for 24 hours before treating them for an additional 24 hours with 10 µM of GM6001, a broad-spectrum MMP inhibitor, and then conducted a scratch assay. We extended the MMP inhibitor treatment during the 8 hours scratch assay. The results in Figure 7 show that the use of GM6001 significantly inhibited the migration for the two patient fibroblasts included in this study, while no effect was obtained for the control cells. These findings strongly suggest that the secretion of MMPs by the cells is directly related to the upregulated fibroblast migration in GD.

**Figure 7.**
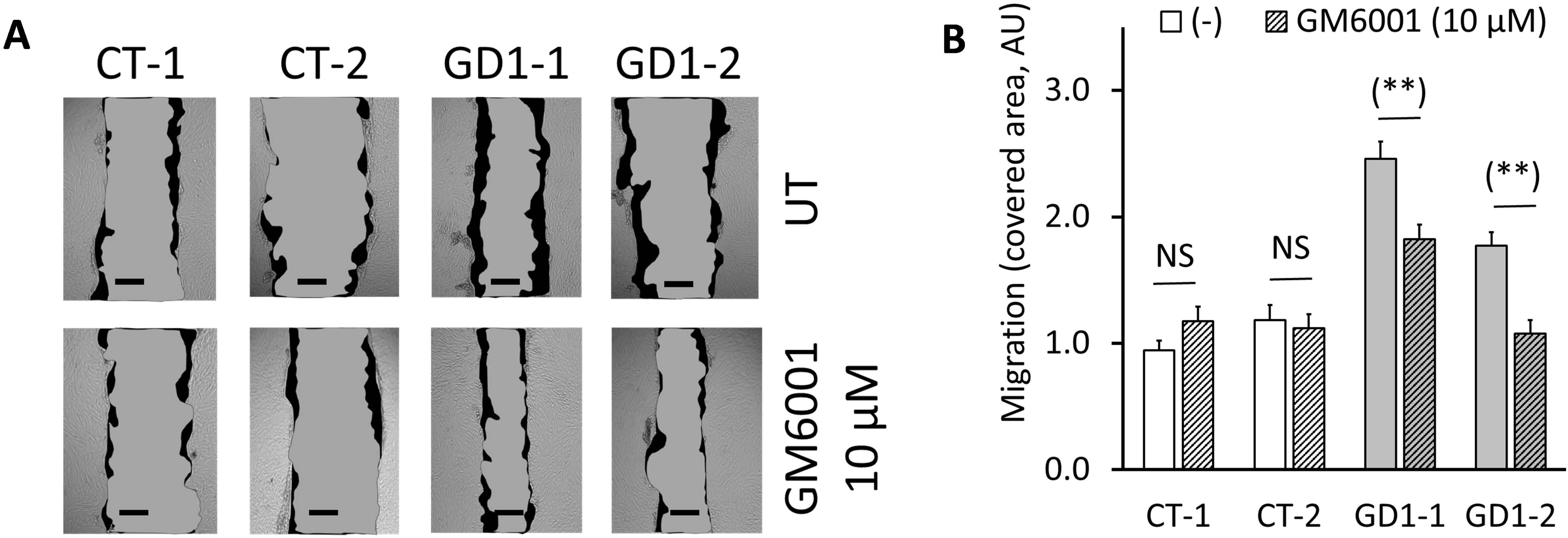
MMPs broad-spectrum inhibitor GM6001, inhibits migration of patient fibroblasts. **(A)** Cells were plated and after 24 hours were treated with 10 µM GM6001. Treatment was extended for 24 hours and during the migration assay. Cell migration was assessed using the wound healing scratch assay for 8 hours. Scale bar = 50 µm. Representative images were selected from four independent experiments, using two CT fibroblasts and two GD1 patient fibroblasts. **(B)** Covered area (dark, in Arbitrary Units) was calculated using ImageJ software. Significance was measured using a t-test and values were compared versus CT fibroblasts (** p<0.01).

## Discussion

Human dermal fibroblasts have been studied as the available cell model of GD, in order to elucidate potential molecular mechanisms of the disease [8,9,11]. Primary skin fibroblasts have been reported with intracellular inclusions (lysosomal-like vacuoles), impaired ADAMTSL2 or FBN1 secretion, reduced and poorly organized ECM microfibrils, and dysregulation of the TGF-β pathway [8–11,48]. Nevertheless, not all GD fibroblasts share similar cellular phenotypes, and the exact mechanism by which variants in *ADAMTSL2* or *FBN1* genes caused this, has not been described yet. Primary dermal fibroblast from five GD patients and three healthy asymptomatic individuals used as controls, were included in this study. No changes in proliferation or cell morphology were observed in the patient’s fibroblasts compared to control cells. However, the amount of ECM fibers and their organization showed notable differences between control and patient fibroblasts; confirming that the deposition of the ECM plays a key role in the pathogenesis of GD, as reported previously [8,9,11]. ECM proteins such as ADAMTSL2, FBN1, and fibronectin were reduced in all patient fibroblasts. Interestingly, FBN1 secretion was affected in fibroblasts with ADAMTSL2 mutations and vice versa. These results suggest a possible interaction between ADAMTSL2 and FBN1 during their synthesis, secretion, and/or incorporation into the ECM; but further studies are required to elucidate the mechanism in their interactions.

The most notable difference between all fibroblasts was the enhanced cell migration of all patient cells. While cell migration is vital in diverse biological processes like tissue repair, development, and the immune response, its dysregulation has been reported in numerous diseases such as cancer, fibrosis and joint diseases [49–51]. Heart and respiratory problems are among the most common and serious complications of GD and the focus for future treatments. Cell migration plays a critical role in heart development, including the migration of cardiac neural crest cells, myocardial precursors, and epicardial cells [52–54]. In addition, cell migration is also a crucial process not only for lung development and growth but also for regeneration after lung injuries. In particular, airway epithelial cell migration, alveologenesis, and smooth muscle cell differentiation have been described as key points for the integrity of the respiratory tissue. They all rely on the interaction of cells with their surrounding ECM molecules [55–59]. Not surprisingly, *ADAMTSL2-KO* mouse models have been reported with very high lethality at birth with severe bronchial epithelial dysplasia and abnormal glycogen-rich inclusions in the bronchial epithelium [18]. Further analysis should explore cell migration during heart and lung development using our ADAMTSL2 mouse model.

Fibroblast migration requires multiple steps for the cells, from extending one edge to form protrusions like lamellipodia or filopodia, adhesion or attachment to the ECM, movement of all cellular contents, and finally a release and retraction of the cell at the trailing edge [60,61]. Particularly important, the ECM is considered a necessary structural support that provides the substrate for attachment and multiple growth factors and other active molecules that can regulate cell migration [62–64]. More recently, Sarkar et al. reported that integrin ligands in the ECM can keep the cells stationary and the depletion of these ligands can spontaneously initiate cell migration [65]. GD fibroblasts showed reduced ECM and faster migration. Surprisingly, growing GD fibroblasts on rich ECM did not affect their migration, suggesting that the high density of integrin ligands cannot inhibit cell migration directly. Nevertheless, we cannot rule out the possibility of a poor interaction of cell integrins and the exogenous ECM ligands.

Cell migration can be triggered by multiple internal and external signals. We found that patient cells overexpressed multiple MMPs. These proteases are key regulators of cell migration by degrading the ECM, which is the substrate for attachment, and by modifying the cell adhesion molecules [66–69]. Specifically, MMP-1 and MMP-14 were significantly up-regulated in patient fibroblasts. To support our hypothesis, we demonstrated that GM6001, a broad-range MMP inhibitor, significantly reduced the patient fibroblast migration. As MMP inhibitors do not target specific MMPs, showing insufficient specificity, we were unable to verify the individual effect of MMP-1 or MMP-14 on fibroblast migration.

Originally, MMPs were identified as proteases primarily responsible for the remodeling of the extracellular matrix with specific distribution among different tissues [36]. We studied human dermal fibroblasts *in-vitro*, and we cannot extrapolate our results for other tissues affected in GD patients. Nevertheless, multiple publications have been focused on MMP profiles from tissues like the heart and the lungs. MMPs are not only key players in heart and lung development and remodeling; but also, they have been reported with critical roles in cardiovascular and pulmonary diseases [70–75]. MMP-1 and MMP-14 are both associated with atherosclerosis and MMP-1 was found increased in heart failure patients and associated with arterial hypertension [71, 76]. Similarly, MMP-1 and MMP-14 are increased in idiopathic pulmonary fibrosis, they are involved in chronic obstructive pulmonary disease (COPD), and they can be induced by smoking and are associated with early-onset of lung cancer [77–81].

We finally confirmed our findings using mouse dermal fibroblasts from a GD mouse model we recently reported, in which the severity of cardiovascular and respiratory dysfunction correlated with impaired secretion of *ADAMTSL2* variants [40]. These mouse dermal fibroblasts showed similar enhanced cell migration when compared with control cells and strong up-regulation of MMP-1 and MMP-14. Additionally, we confirmed the dysregulation of MMPs in the lung and heart of *ADAMTSL2-KO* mice, especially the strongly up-regulated MMP1 and MMP14 in these tissues. These results not only validated our findings in human dermal fibroblast but also confirmed our animal model as a valuable tool for future studies.

In conclusion, when comparing GD fibroblasts with control cells, most of their characteristics resulted in subtle differences except the faster migration of GD fibroblast via up-regulation of MMPs. A better understanding of the expression profile of MMPs and their function during tissue development and remodeling, especially in heart and lung, and their regulation during associated diseases could provide future opportunities and therapeutic options for GD patients.

## Material and Methods

### Study approval

All human and animal studies were reviewed and approved by the University of Miami Institutional Review Board (IRB) and by the University of Miami Institutional Animal Care and Use Committee (AICUC), according to the National Institutes of Health guidelines (NIH), and all patients were evaluated at the University of Miami or by collaborators. Written informed consents for the testing and sharing of the research data were obtained from all patients or parents, including the use of the isolated primary fibroblasts.

### Cell culture experiments

All dermal fibroblasts were isolated from skin biopsies, following the explant technique, except GD006 (GD1-2) which was kindly provided by the Bristol Genetics Laboratory at North Bristol NHS Trust, Severn Pathology, Southmead Hospital, UK. Briefly, tissues were cut into small explants and incubated at 37°C and 5% CO2 atmosphere, using complete Dulbecco’s Modified Eagle Medium (DMEM) high glucose, pyruvate, 20% FBS, 100 U/mL of penicillin, 100 μg/mL of streptomycin, and 200 μg/mL of Primocin. The medium was carefully changed every day until cells migrated from the explant and grew as a monolayer. Dermal fibroblasts were cryopreserved as passage P2. Thawed fibroblasts for experiments were cultured in T75 flasks and used in a range from P4-P10 passages, although they were all able to further expand for more than 20 passages, without visible changes in size, morphology, or capacity to proliferate. All experiments were completed using Dulbecco’s Modified Eagle Medium (DMEM) high glucose, pyruvate, 10% FBS, 100 U/mL of penicillin, 100 μg/mL of streptomycin, unless a different medium is specified. HDFn (CT) cells are commercially available (Invitrogen, C-004-5C).

### Cell Proliferation Assay

Fibroblast proliferation was assessed by evaluating the growth rates for 10 days. Cells were plated at an initial density of 1.5 ×10^4^ and 7.5×10^4^ cells per well, in 96-well plates and allowed to attach and grow during 10 days at 37°C and 5% CO2 atmosphere. Cell proliferation was evaluated at 1, 2, 4, 7, and 10 days, using the Lactate Dehydrogenase (LDH) activity assay (Cytotoxicity Detection Kit PLUS, Roche, 04744926001), a colorimetric test that measures the amount of LDH with the reduction of a yellow tetrazolium salt into a red formazan dye. Cell proliferation was determined as total LDH content per well after complete lysis of cells. LDH content at time 0 was considered 100% and the percentage of total LDH per well calculated for each condition and time point. Experiments were performed in triplicates and five independent experiments were averaged.

### Scratch Assay

For the wound healing assay, 5×10^5^ cells were plated per well, in 12-well plates. Cells were allowed to attach and grow for 48 hours at 37°C and 5% CO2 atmosphere, for a 100% monolayer confluency. Gentle scratch using 200 µL tips were performed in each well. Cells were washed with PBS and fresh complete Dulbecco’s Modified Eagle Medium was added. Images were captured at 0, 8 and 24 hours post scratch using a camera attached to the microscope at 4x. Four different fields were used per replicate, for a total of 8 fields per cell, per experiment (two replicates per sample per condition). The wound area was measured by tracing the edges of the wound, using ImageJ software [82]. Five independent experiments were averaged. A similar protocol was followed for the migration of fibroblasts on top of a mature ECM. HDFn cells were cultured for 7 days, and the mature ECM was isolated following the Ammonium hydroxide (NH4OH) / Triton-X100 procedure [83].

### Protein lysates

Protein lysates were prepared using four different approaches depending on the experiment goal.

1. Cell lysates (CL) were prepared using 1x RIPA lysis buffer (Millipore-Sigma, R0278) containing 50 mM Tris-HCl, pH 8.0, with 150 mM sodium chloride, 1.0% Igepal CA-630 (NP-40), 0.5% sodium deoxycholate, and 0.1% sodium dodecyl sulfate (SDS), plus a 1x protease inhibitor mixture (C0mplete™ Protease Inhibitor Cocktail, Roche, 11697498001). RIPA was added directly to the cells and then the cells were scraped and incubated on ice for 15 minutes. Cell lysates were cleared at maximum speed (14000 rpm) for 5 minutes.
2. Intracellular lysates (IC) were prepared by adding RIPA buffer to trypsinized cells and incubated on ice for 15 minutes. Trypsin-EDTA (0.25%, Gibco 25200056) was added to the cells and incubated at 37°C for 5 minutes. Trypsin was inhibited using complete medium DMEM(10% FBS) and cells were washed with PBS. Intracellular lysates were cleared at maximum speed (14000 rpm) for 5 minutes. Protein concentration was determined using a Bradford colorimetric assay kit (Bio-Rad Protein Assay Kit-II Bio-Rad, 500-0002).
3. Total lysates (Total) were prepared adding 2x Laemmli buffer (Bio-Rad, 1610747), directly to the cells in well, previously washed with PBS.
4. Extracellular matrix (ECM) protein lysates were prepared using the Ammonium hydroxide (NH4OH) / Triton-X100 protocol [83]. Briefly, cells were washed with PBS and removed from the plate with 20 mM NH4OH, 0.05% Triton-X100 in PBS for 15 minutes at room temperature. The remaining ECM proteins attached to the plate were washed with PBS and dissolve in 2x Laemmli buffer as 138.9 mM Tris-HCl, pH 6.8, 22.2% (v/v) glycerol, 2.2% LDS and 0,01% bromophenol blue containing 10% 2-Mercaptoethanol (Sigma).

### Tissue lysates

lungs and hearts from WT and ADAMTSL2-KO neonate mice (P1) were cut in pieces around 1-2 mm^2^ and manually homogenized using a Cole-Parmer® Motorized Pestle Mixer. Then, 1x RIPA lysis buffer was added in a ratio of 1 mL of lysis buffer per 10 mg of tissue and briefly homogenized again. Tissue lysates were extracted in a laboratory shaker at 4oC for 30 minutes and finally cleared at maximum speed (14000 rpm) for 5 minutes.

### Western Blot Analysis

Ten to twenty µg of proteins were subjected to SDS-polyacrylamide gel electrophoresis (SDS-PAGE), using the appropriate gel, and transferred to 0.22 µm PVDF membrane (Bio-Rad, 1620112). After blocking with 5% BSA in Tris Buffered Saline (TBS-T20, 25mM Tris, 140 mM Sodium Chloride, and 3.0 mM Potassium Chloride, and containing 0.1% Tween-20); the membranes were incubated with the corresponding primary antibody overnight at 4°C (Table.1S). We used a polyclonal antibody against the C-terminus of ADAMTSL2 (C3) from GeneTex and it is the best available antibody. Primary antibodies were detected by horseradish peroxidase conjugated secondary antibody and the immunocomplexes were visualized by chemiluminescence (Super Signal™ West, Thermo-Fisher Scientific, 34577 and 34096). The membranes were stripped using Restore™ PLUS Western Blot Stripping Buffer (Thermo Scientific, 46430) at room temperature, for 15 minutes, extensively washed with TBS-T20, blocked with 5% BSA in TBS-T20, and reprobed with the corresponding antibodies following a similar protocol.

### Whole transcriptome sequencing (RNA-seq)

Total RNA was extracted from primary human fibroblast, at P4. The cells were grown in DMEM 10% FBS, but the serum was removed overnight the day before the collection. The RNA was extracted using the RNeasy Mini Kit (Qiagen). A Bioanalyzer 2000 was used to measure the quality of RNA. All samples’ RNA integrity numbers (RIN) were above 9. Whole transcriptome sequencing (RNA-seq) was conducted at the Sequencing Core of John P. Hussman Institute of Human Genomics at the University of Miami using the TruSeq Stranded Total RNA Library Prep Kit from Illumina (San Diego, CA). Briefly, after ribosomal RNA (rRNA) was depleted, sequencing libraries were ligated with standard Illumina adaptors and subsequently sequenced on a Hiseq2000 sequencing system (100 bp single end reads, Illumina, San Diego, CA, USA). Trimming was performed, and sequence reads were aligned to the human transcriptome (GRCh38/hg38 assembly from the Genome Reference Consortium) and quantified using the STAR aligner [84]. All samples had uniquely aligned reads between 35,607,220 and 51,912,532. Read data was run through quality control metrics using MultiQC [85]. Statistical significance was first examined with three alternative differential expression calculators: edgeR, DESeq2, and Bayseq [86–88]. For visualization and further analysis iDEP software was used [89]. The program calculates CPM values from raw counts. To reduce false positive a minimum of 5 CPM per library was used. Data was transformed, normalized for clustering with rlog (transformation implemented in DESeq2). Of 60,656 genes from the data set, 11,147 genes passed the filter. A supervised Hierarchical and K-Means clustering with heatmap was generated to have a specific view of the expression of metalloprotease family of genes.

### MMP activity

Active and total MMP activity was determined using the green fluorometric MMP Activity Assay Kit (Abcam, ab112146) following manufacturer instructions. Briefly, samples (conditioned media) were incubated with equal volume of Assay Buffer (for active MMPs) or 2 mM 4-Aminophenylmercuric acetate (APMA) working solution (for Total MMPs) at 37°C for 2 hours. In a 96-well plate, combine 50 µL of each sample with 50 µL of MMP Green Substrate working solution. Mix well all samples and monitor the fluorescence intensity at 0, 15, 30, 60 and 90 minutes at room temperature, with a fluorescence plate reader at Ex/Em = 490/525 nm. Fluorescence from all samples were subtracted with the reading for control DMEM+10%FBS used to obtain the conditioned media. Relative fluorescence units (RFUs) were averaged from at least three independent experiments.

### MMPAntibody Array

MMPs were detected using the Human MMP Antibody Array (Abcam, ab134004) following the manufacturer instructions. Briefly, membranes were blocked with the Blocking buffer at room temperature for 30 minutes and incubated with the conditioned media overnight at 4°C. Membranes were washed and incubated with the Biotin-Conjugated Anti-MMPs for 2 hours at room temperature. Membranes were washed again and incubated with the HRP-Conjugated Streptavidin for 2 hours at room temperature. Membranes were washed again and incubated with the Detection buffer at room temperature for 2 minutes to visualize the immunocomplexes by chemiluminescence. Spot intensities were determined using ImageJ software.

### Statistics

Data are expressed as mean ± SEM and all graphs were performed in Microsoft Excel. The number of replicates for experiments are indicated in each figure and statistical differences were tested using the Student t-test when comparing a group with the corresponding control. P-values < 0.05 were considered statistically significant as * p<0.05, ** p<0.01, and *** p<0.001.

## Author contributions

AAM designed and conducted experiments, acquired data, analyzed data, and wrote the manuscript. VC, GW and KW analyzed data and reviewed the manuscript. LP, SS, LS, LW, JMN, JES and AB recruited patients and reviews the manuscript. MT designed the research study, analyzed data, and reviewed the manuscript.

## Data availability

The datasets generated and/or analyzed during the current study are available in the ClinVar repository. Submission IDs for reported gene variants include: ADAMTSL2 c.182G>A as SUB15022370, ADAMTSL2 c.493G>A as SUB15022407, ADAMTSL2 c.542T>C as SUB15022419, ADAMTSL2 c.707C>T as SUB15022429 and FBN1 c.5284G>A as SUB15022437, FBN1 c.5183C>T as SUB15022369 and FBN1 c.5243G>C as SUB15022471.

## Acknowledgments

We are immensely grateful to the patients for their participation in this study. This study was supported by a gift from the Al Rashid Family.

## Supplemental Figures and Tables

**Figure 1S.**
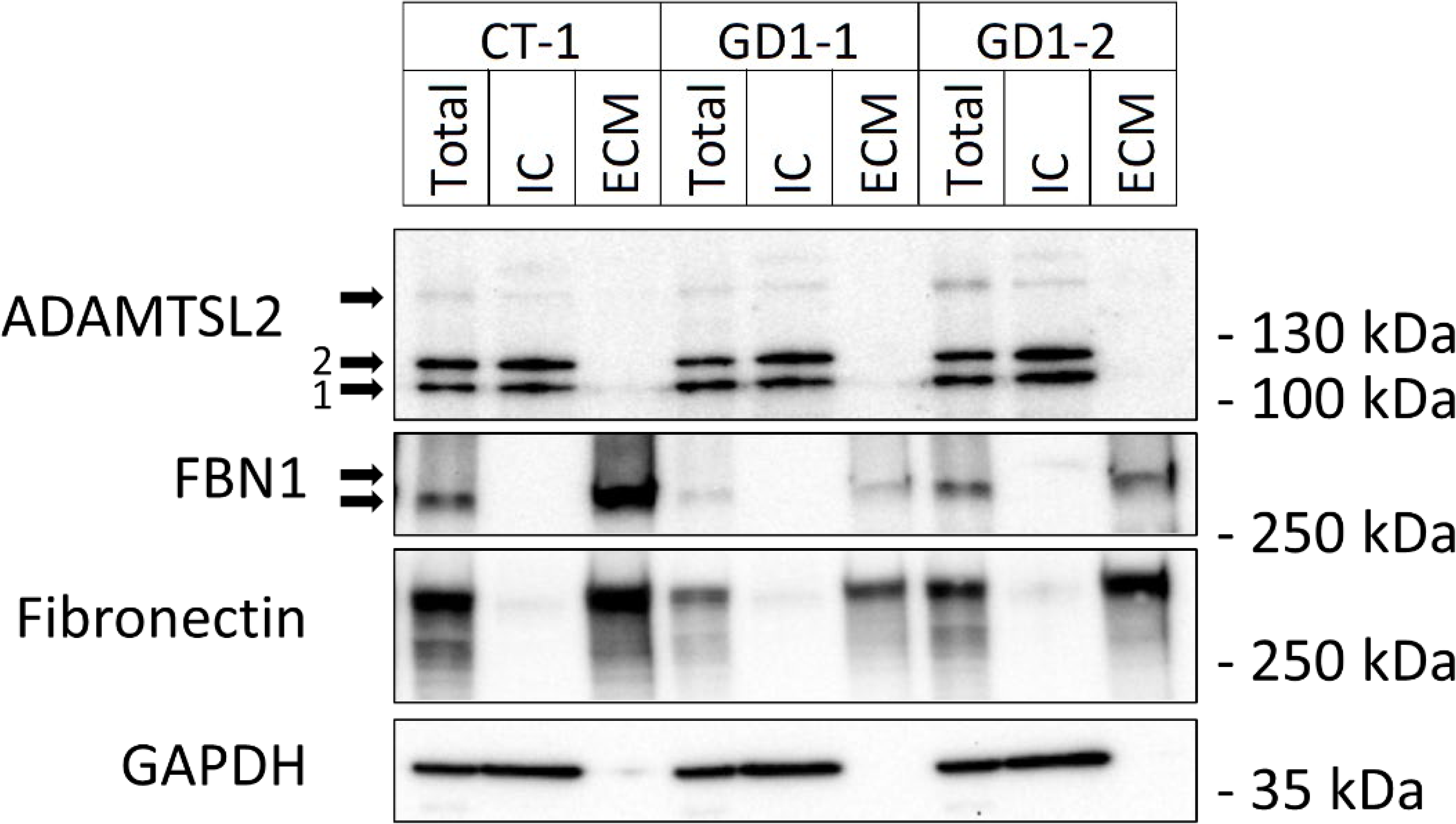
Patient fibroblasts show impaired ECM protein accumulation. Cell lysates were prepared for one CT fibroblast and two GD1 patient’s cells. Total lysates (Total), intracellular lysates (IC) and extracellular matrix lysates (ECM) were analyzed by WB, using GAPDH and Fibronectin as IC and ECM controls, respectively. ADAMTSL2 was detected mainly in the IC compartment while FBN1 and Fibronectin proteins were detected mainly in the ECM space.

**Figure 2S.**
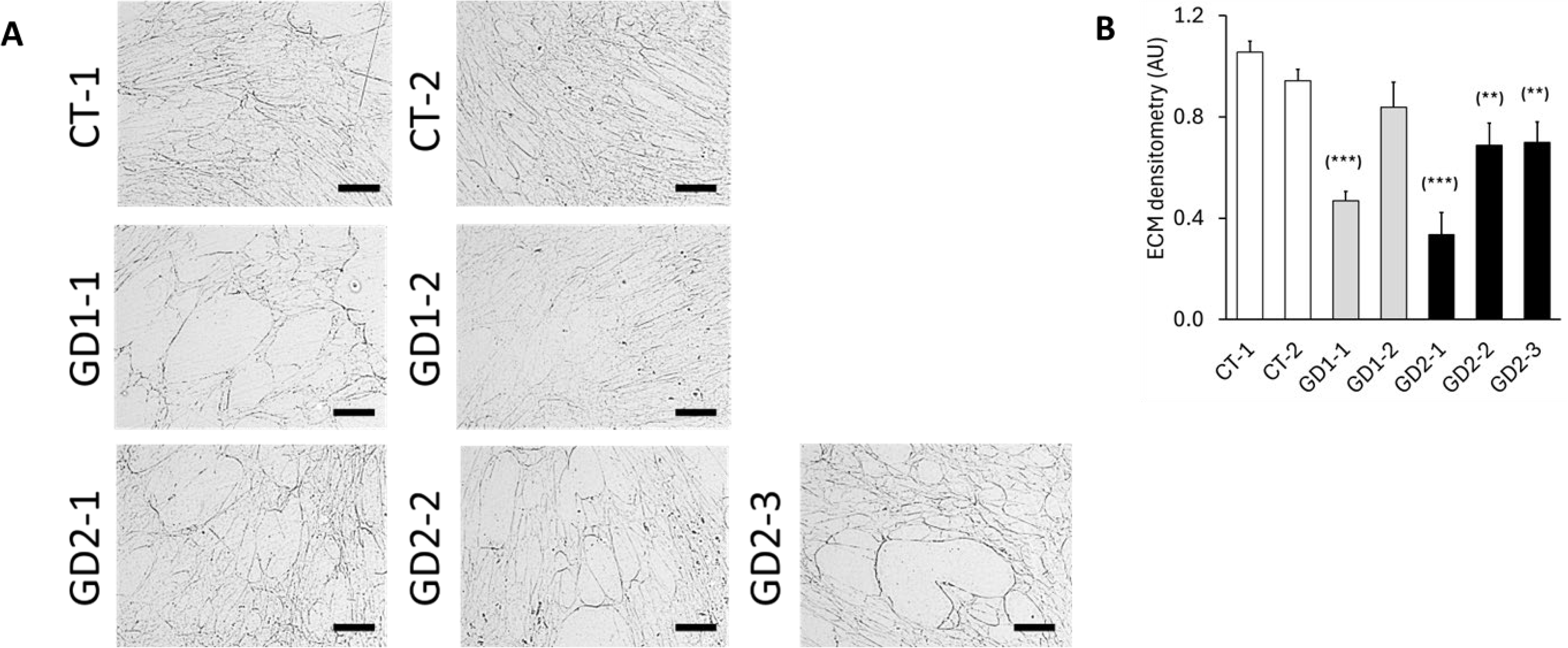
Patient fibroblasts show impaired ECM protein accumulation. Cells were grown for seven days. ECMs were isolated using the NH4OH / Triton-X100 protocol. Representative images were selected from 5 independent experiments. Scale bar = 200 µm.

**Figure 3S.**
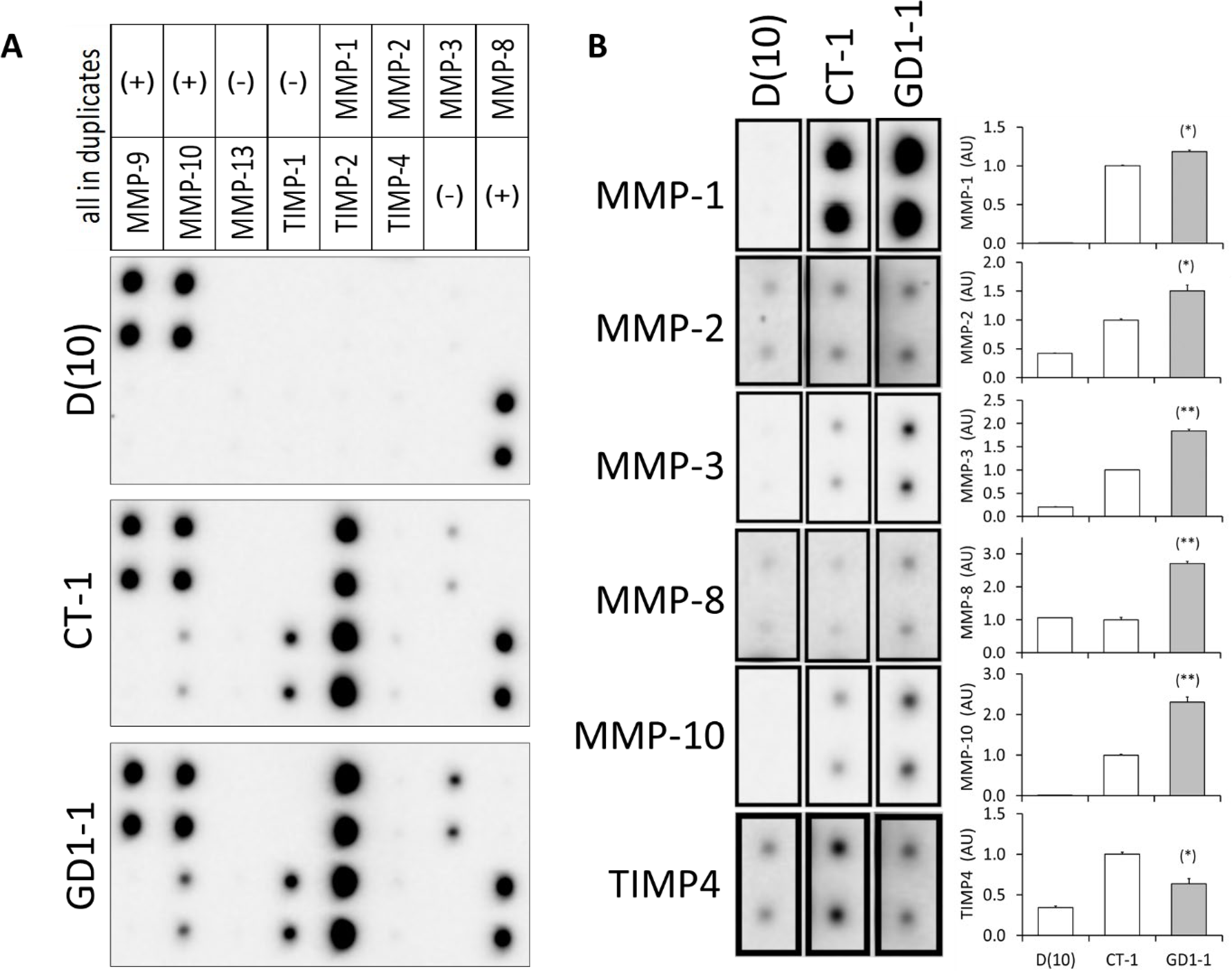
Patient fibroblasts show higher secretion of MMPs. (A) Multiple MMPs were simultaneously detected in the conditioned medium (10 days) of CT and GD1 fibroblasts, using an antibody array (Human MMP Antibody Array). Medium (DMEM+10% FBS) was used as negative control. (B) Densitometry analysis was performed using ImageJ software, for up-regulated or down-regulated proteins. Significance was measured using a t-test and values were compared versus CT fibroblasts (* p<0.05, and ** p<0.01).

**Table 1S.** Antibodies used in this study.

| **Antibody** | **Company** | **Catalog** |
| --- | --- | --- |
| **ADAMTSL2** | GeneTex | GTX102069 |
| **Fibronectin** | Sigma-Millipore | F0916 |
| **FBN1** | Sigma-Millipore | PA5-99225 |
| **Col1A1** | Novus Biologicals | NB600-408 |
| **Col3A1** | Novus Biologicals | NB600-594 |
| **GAPDH** | Cell Signaling Technology | #5174 |
| **MMP-1** | Cell Signaling Technology | #54376 |
| **MMP-2** | Cell Signaling Technology | #40994 |
| **MMP-7** | Cell Signaling Technology | #71031 |
| **MMP-14** | Cell Signaling Technology | #26424 |
| **TIMP1** | Cell Signaling Technology | #8946 |
| **TIMP2** | Cell Signaling Technology | #5738 |
| **MMP-1 (Mouse)** | Invitrogen | PA527210 |
| **MMP-14 (Mouse)** | Invitrogen | PA513183 |

